# Endogenous pathology in tauopathy mice progresses via brain networks

**DOI:** 10.1101/2023.05.23.541792

**Authors:** Denise M.O. Ramirez, Jennifer D. Whitesell, Nikhil Bhagwat, Talitha L. Thomas, Apoorva D. Ajay, Ariana Nawaby, Benoît Delatour, Sylvie Bay, Pierre LaFaye, Joseph E. Knox, Julie A. Harris, Julian P. Meeks, Marc I. Diamond

## Abstract

Neurodegenerative tauopathies are hypothesized to propagate via brain networks. This is uncertain because we have lacked precise network resolution of pathology. We therefore developed whole-brain staining methods with anti-p-tau nanobodies and imaged in 3D PS19 tauopathy mice, which have pan-neuronal expression of full-length human tau containing the P301S mutation. We analyzed patterns of p-tau deposition across established brain networks at multiple ages, testing the relationship between structural connectivity and patterns of progressive pathology. We identified core regions with early tau deposition, and used network propagation modeling to determine the link between tau pathology and connectivity strength. We discovered a bias towards retrograde network-based propagation of tau. This novel approach establishes a fundamental role for brain networks in tau propagation, with implications for human disease.

**One-Sentence Summary:** Novel whole brain imaging of p-tau deposition reveals retrograde-dominant network propagation in a tauopathy mouse model.

## Introduction

Neurodegenerative tauopathies are characterized by progressive neuronal and glial accumulation in amyloid assemblies of tau protein (*1*). Several lines of evidence indicate tau propagates pathology among neurons. Tau assemblies enter cells to trigger intracellular aggregation (*2, 3*), and inoculation of mice with tauopathy homogenates produces local pathology that spreads beyond the injection site (*4*). Prion protein (PrP) inoculation also propagates pathology in neuroanatomical networks (*5*). Human studies based on functional connectivity and tractography have suggested brain networks could play a role in tauopathy progression, consistent with the prion hypothesis (*6, 7*). Tau also forms strains, which are defined assembly conformations that propagate faithfully *in vivo*, creating unique transmissible pathologies (*8, 9*). Finally, immunotherapies targeting extracellular tau reduce pathology in mice (*10*). However, despite being a core feature of the prion model, this hypothesis remains uncertain.

In humans, MRI-based studies using fMRI and tractography cannot precisely map pathology at the cellular level. Instead, they rely on atrophy patterns or relatively low-resolution PET imaging of tau deposition, and histopathological analysis of end-stage disease (*7*). Second, while inoculation in rodents has been widely used by our lab and others to induce focal pathology that spreads from a defined brain region (*1, 4, 8, 9, 11-13*), these methods induce brain trauma, cannot exclude trafficking of inoculated seeds within a network, and do not reflect the spontaneous development of pathology that occurs in patients. Finally, traditional methods of assessing tau pathology via immunostaining of serial (2D) brain sections pose a barrier to complete, unbiased assessment of pathological progression. Advances in image registration and analysis of serial sections improve this capacity (*13, 14*), but remain laborious and limited to a subset of anatomical levels. Consequently, the hypothesis of trans-neuronal propagation of endogenous tau pathology, while intriguing, has remained fundamentally untested *in vivo*.

We now report novel methods for labeling, volumetric imaging, and analysis of the progression of spontaneous tau pathology in PS19 mice. We have tested the network model by combining 3D maps of progressive pathology and the mouse structural connectome (*15*), without confounds introduced by inoculation or expression within restricted neuronal populations. Using a fluorescently tagged camelid nanobody (VHH) directed against pathological human tau phosphorylated at residue S422 (VHH-A2-488) (*16*), we determined phosphorylated-tau (p-tau) deposition in intact, uncleared brains of PS19 mice at 3 to 12 months. Despite widespread transgene expression, high-resolution whole-brain images revealed p-tau neuropathology that began spontaneously in a small number of distinct brain regions. Whole brain analysis of tau pathology defined characteristic, non-random patterns of p-tau deposition that progressed over time, with a bias toward retrograde propagation.

## Results

### Development of whole brain p-tau staining and quantification pipeline

Building on previous studies (*16-22*), we created a high-resolution imaging and informatics pipeline to map the distribution of pathological tau across the whole brain in 3-12mo PS19 mice (Fig. 1A). These animals express 1N4R human tau containing a disease-associated mutation (P301S) driven by the pan-neuronal prion promoter (*23*). We used a previously characterized, fluorescently conjugated nanobody directed against human tau (pS422) (VHH-A2-488), which stains Alzheimer’s and P301L tauopathy mouse brain (*24*) comparably to AT8 (*16*), and an anti-p-tau monoclonal antibody widely used to monitor pathology (*25-28*). Because of their smaller size and other inherent features (*29, 30*), we expected VHH-A2-488 to better penetrate and diffuse through large tissue volumes—essential for serial two-photon tomography (STPT). We optimized a protocol for permeabilization, immunolabeling and washing compatible with whole brain staining of p-tau (Fig. 1A).

**Fig. 1.**
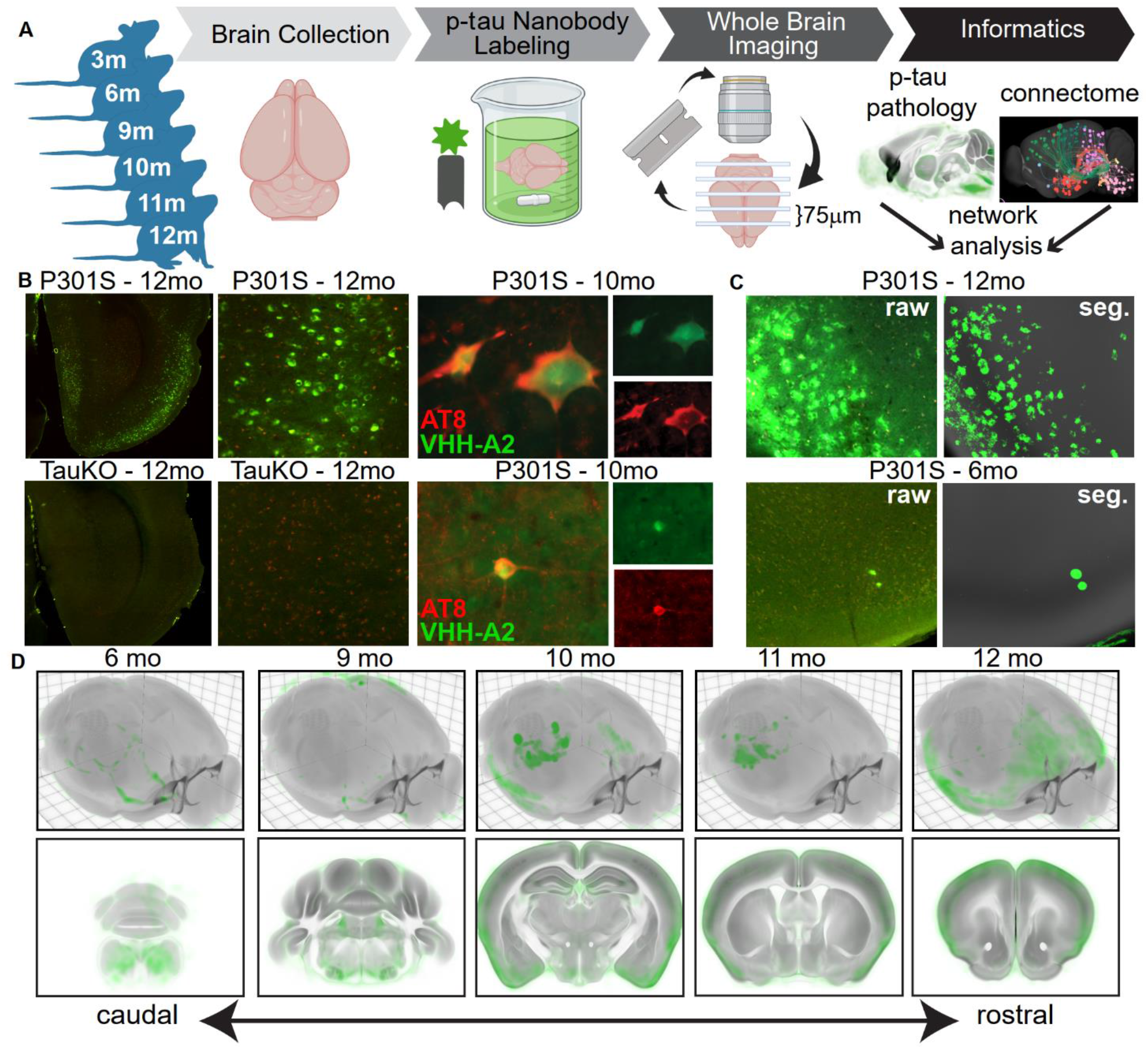
Whole brain imaging and analysis of nanobody-stained p-tau pathology. (A) Schematic of experimental workflow. Twenty-three PS19 mice were sacrificed at ages 3, 6, 9, 10, 11, or 12mo and immunostained with VHH-A2-488. The brains were subjected to STPT, generating a library of 3-dimensional whole-brain images of spontaneous p-tau deposition. A custom informatics workflow incorporating supervised machine learning-based pixel classification and registration into the CCFv3.0 was used to quantify p-tau accumulation across the cohort. Network diffusion modeling was used to compare the patterns of brain-wide spontaneous tau pathology to structural brain networks. (B) Upper left, p-tau staining in a section from entorhinal cortex (ENT) of a 12mo PS19 mouse brain and enlarged to show p-tau neuronal morphology. Lower left, staining of 12mo TauKO brains with VHH-A2-488 did not produce neuronal signal. Green = VHH-A2-488 positive p-tau deposits, Red = tissue autofluorescence. Upper and lower right, co-staining of AT8 (red), a canonical marker for p-tau, and VHH-A2-488 (green) showed consistent examples of double labeled neurons. (C) Image taken from piriform cortex (PIR) of PS19 animals stained with VHH-A2-488 at 12mos (upper panels) or 6mos (lower panels). Left side shows raw fluorescence images (VHH-A2-488 in green) and right side shows the corresponding probability map outputs of the same region (segmented p-tau signal in green and atlas template in grey). (D) Average brain-wide p-tau distribution in PS19 mice in each age group (6, 9, 10, 11, and 12mos). P-tau probabilities are shown in green and the CCF average template in grey. Top, 3D renderings of the whole average brain from a dorsal oblique view. Bottom, Coronal planes of the 12mo average brain spanning the rostro-caudal extent of the brain. Accumulation of p-tau is visible across many brain regions, notably brainstem, hippocampus and cortex, and these regions exhibited a high p-tau burden in 12mo animals. Number of brains averaged for each age group: 6mo, n=5; 9mo, n=3; 10mo, n=6; 11mo, n=5; 12mo, n=4.

STPT images of VHH-A2-488 immunostained brains demonstrated neuronal p-tau staining across brain regions including entorhinal cortex (EC; Fig. 1B), piriform cortex (PIR; Fig. 1C) and anterior hypothalamic nucleus (AHN; Fig. S1). As observed previously (*16*), neurons stained with VHH-A2-488 were sometimes co-labeled with AT8 antibodies (Fig. 1B). We also observed independent populations of VHH-A2-488- and AT8-positive neurons (data not shown) which might be explained by the different tau phospho-epitopes targeted by each antibody. In tau knockout brains (*31*) we observed no neuronal p-tau staining by VHH-A2-488, confirming its specificity (Fig. 1B and Fig. S1). We then developed a machine learning model compatible with variations in background intensity or texture to classify pixels as VHH-A2-positive or -negative (Fig. 1C, Fig. S1). We produced “probability maps” of p-tau that were transformed into the 3D Allen Common Coordinate Framework version 3 (CCFv3) reference atlas (*17, 18, 20, 21*).

We visually inspected the CCFv3-registered p-tau probability maps and observed consistent patterns of tau pathology at all ages (Fig. 1D, Fig. 2, Fig. S2, Fig. S3). 6mo animals exhibited low p-tau staining above background, most consistently in brainstem and other caudal regions (Fig. 1D, Fig. 2A-D, Fig. S2A-D, Fig S3A-C). At 9-10mos, p-tau signal increased in brainstem regions, and low-to-moderate signal appeared in more rostral regions, including hippocampus and isocortex (Fig. 1D, Fig. 2E-L, Fig. S2E-L, Fig. S3D-I). Despite inter-animal variability at each age, a common set of brain structures with p-tau pathology emerged when comparing 6, 9, and 10mo mice, including the pons (P), medulla (MY), hypothalamus (HY), entorhinal cortex (ENT) and piriform cortex (PIR). This suggested p-tau accumulated non-randomly. The 11-12mo animals often had intense p-tau pathology in many brain structures (Fig. 1D, Fig. 2 M-T, Fig. S2M-T, Fig. S3J-O). P-tau positive structures in 11-12mo brains largely overlapped those at 9-10mo. The tau burden in specific regions was not necessarily higher than in the younger animals, possibly owing to degenerative changes in the older brains. The 11-12mo brains also exhibited pathology in anterior regions such as somatosensory (SS) and motor (MO) cortices, suggesting the network of tau-susceptible structures expanded with age. The general pattern of p-tau accumulation matched recent reports of brainwide p-tau accumulation in the P301L tauopathy model (*12, 24*). A subset of brains showed especially strong p-tau pathology in the isocortex (Movie S1). Staining for human tau (with HJ8.5 antibody (*3, 10*)) indicated that p-tau deposition did not correlate with total tau expression (Fig. S4). Together, these data strongly suggested that p-tau accumulation in the PS19 mouse model was ordered and progressive in nature, and could not be attributed simply to transgene expression.

**Fig. 2.**
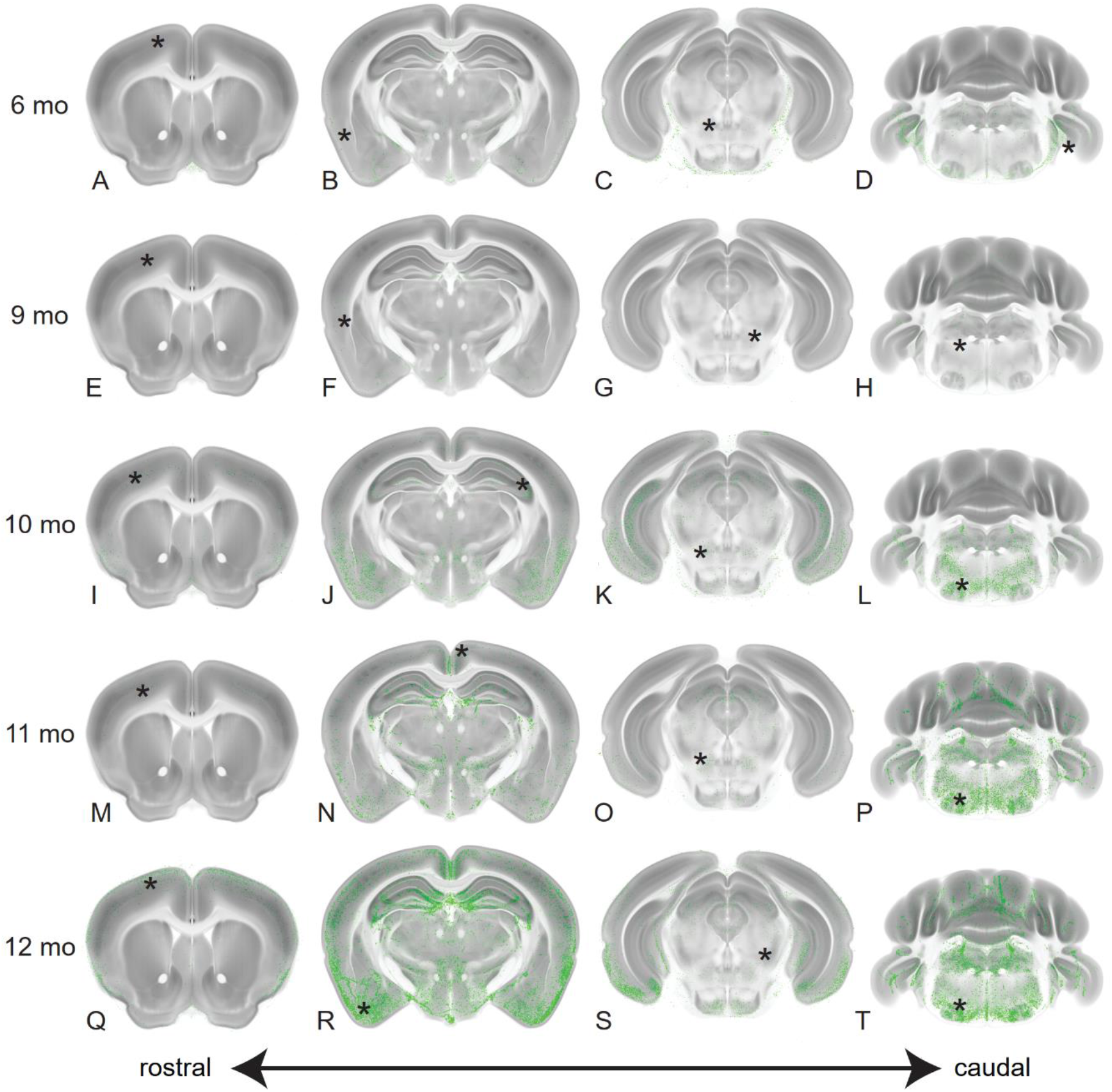
P-tau pathology initiates caudally and progresses rostrally. We show maximum intensity projections (MIP) of p-tau probability maps in each age group at selected anatomical levels. Four coronal sections spanning the rostro-caudal extent of the brain at the levels of the anterior cortex, dorsal hippocampus, midbrain and cerebellum are shown for brains of the indicated ages (6mo = Panels A-D; 9mo = Panels E-H; 10mo = Panels I-L; 11mo = Panels M-P; 12mo = Panels Q-T). P-tau probabilities are shown in green and the CCF average template in grey. The total numbers of brains in the dataset in each age group (6mo, n= 5; 9mo, n= 3, 10mo, n= 6; 11mo, n= 5; 12mo, n= 4) were used to generate MIP images of each physical section. Progressive accumulation of p-tau was visible across many brain regions, notably brainstem, hippocampus and ventral cortical areas including entorhinal (ENT), cortical amygdala (COA) and piriform (PIR) regions. Asterisks on each section indicate the location of the corresponding enlarged images shown in Fig. S2.

### Nonrandom patterns of tau pathology emerge across animals and ages

In humans, neuropathological heterogeneity has been proposed to reflect variable onset times, anatomy, and rates of disease progression (*6*). We investigated whether p-tau staining patterns represented mild-to-severe variants of pathology in a consistent set of brain regions, or whether they were more random. To approach this question, we used cluster analysis on region-normalized p-tau signals across 23 brains and 315 brain regions annotated in the Allen CCFv3. These “summary structures” covered the entire brain (Fig. 3) (*21, 32*). This revealed pathological patterns related to, but spanning, age groups (Fig. 3A). We observed no obvious asymmetries between hemispheres in p-tau staining at any age (Pearson’s correlation coefficient 0.9226, p < 1e-12). Cluster analysis confirmed initial visual observations of p-tau pathology in specific brain structures. P-tau signal was generally highest in the pons (P), medulla (MY), and select structures in the cerebellum (CB) and hypothalamus (HY). The cluster with lowest overall p-tau intensity (Fig. 3A, “C1”) was found in five 6mo brains, when consistent pathology emerges in the PS19 model (*3, 23*). A second cluster with relatively mild caudal staining contained four older (10-11mo) brains (Fig. 3A, “C2”), indicating some animals were resistant to pathological progression. A third cluster of four 9-10mo brains had pronounced caudal staining, including P, MY, and CB (Fig. 3A, “C3”). The fourth cluster included five 10-12mo brains with intense p-tau staining in caudal regions and increased pathology in midbrain (MB), olfactory areas (OLF), and the hippocampal formation (HPF)(Fig. 3A, “C4”). Finally, three 10-12mo brains clustered together that had a distinct pattern of pathology with pronounced staining in isocortex, OLF, HPF, and only moderate staining in P, MY, and CB (Fig. 3A, “C5”). Overall, we found highly nonrandom patterns of p-tau pathology, with C1-C4 appearing to be instances of pathological progression across one distinct subcortical set of regions (“Subcortical”), and C5 appearing to be a late stage of progression across a separate, distinct set of cortical regions (“Cortical”). Remarkably, excepting a few caudal structures, brain-wide patterns of neuropathology in the Cortical and Subcortical groups were almost entirely nonoverlapping (Fig. 3A, 3B, Movie S1). With the caveat that our study was not powered to assess sex differences as a primary outcome, we found no obvious effects of sex on either pathological burden or presentation of the subcortical vs. cortical pattern of pathology (Table S1).

**Fig. 3.**
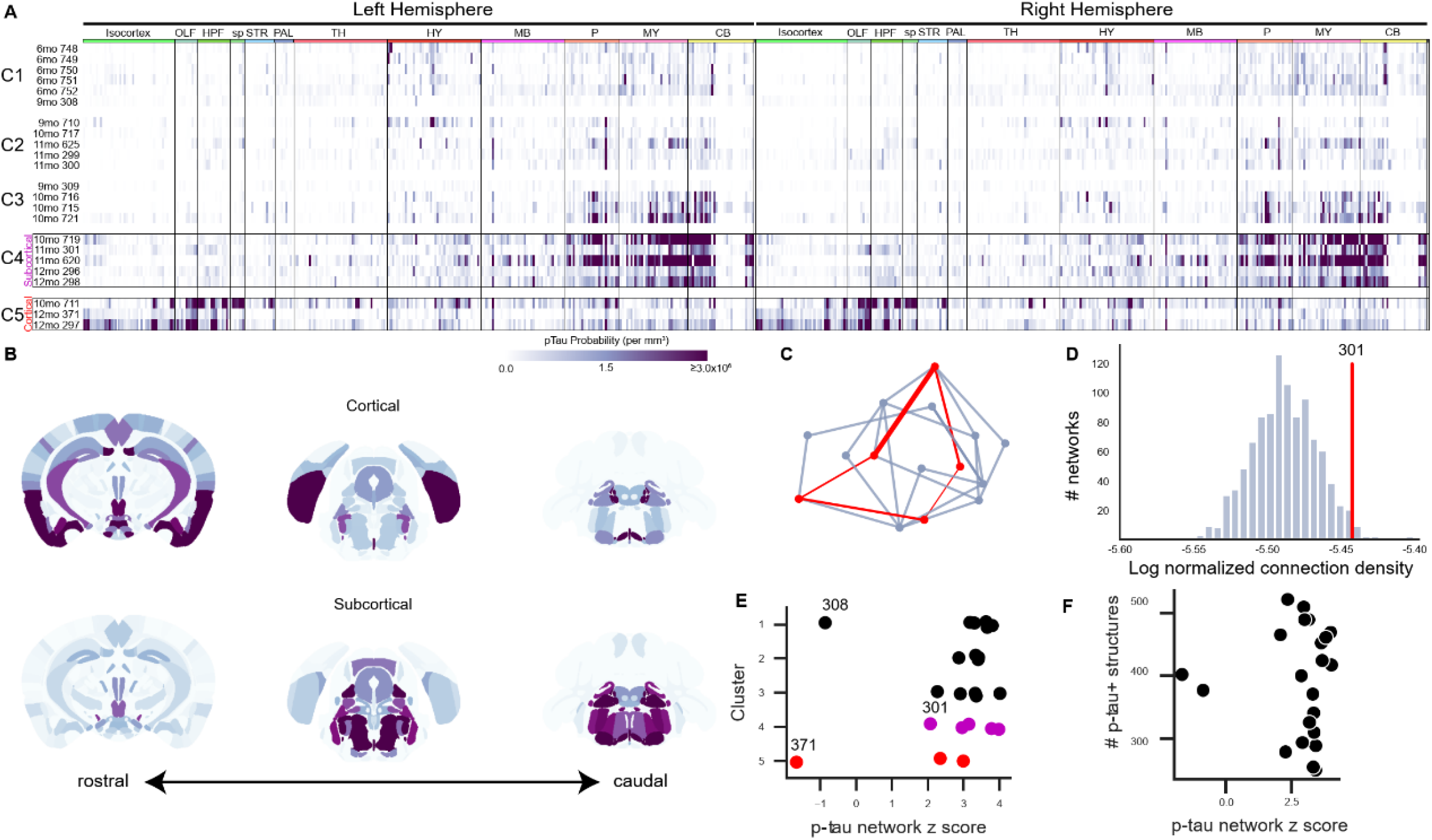
Patterns of spontaneous p-tau pathology match structural connectivity. (A) Raster plots showing automated quantification of region-normalized p-tau probability levels in each of the 630 structures annotated in the Allen Institute CCFv3. Displayed values were truncated at 3×10^6^ to visualize lower-intensity regions. Clustering based on brain-wide patterns of pathology identified progressive accumulation of p-tau in a predominantly subcortical pattern (C1-C4), but a subset of three brains exhibited an alternate pattern with heavier pathology in cortical regions (C5). Both “Subcortical” (C4) and “Cortical” (C5) clusters of brains with high p-tau burdens were composed of aged mice (10-12mo). Labels across the top indicate major brain divisions. Abbreviations: OLF: Olfactory areas; HPF: Hippocampal formation; sp: Cortical subplate; STR: Striatum; PAL: Pallidum; TH: Thalamus; HY: Hypothalamus; MB: Midbrain; P: Pons; MY: Medulla; CB: Cerebellum. (B) Maps showing the spatial pattern of p-tau pathology by annotated brain region. Median values are shown for n=3 brains with pronounced Cortical pathology patterns and n=5 brains with Subcortical pathology (indicated by boxes in A). Coronal sections at the levels of the hippocampus (left), midbrain (center), and brainstem (right) are shown for each pattern. Heatmap is the same as A. (C) Schematic of method used to evaluate whether p-tau levels were related to connectivity. For each brain, the set of regions (nodes) that contained p-tau signal (red points) were identified and the sum of the anatomical connection strength between regions was calculated by summing all the edges in this graph (red lines). Inter-regional connection strength values came from the regionalized voxel model in Knox et al. (*33*). The strengths of all the connections in this network were summed and compared to results of 1000 alternative networks drawn from the same brain-wide connectivity matrix and containing a similar distribution of inter-regional distances (blue-gray lines). (D) Histogram of the total connection strength of 1000 alternative networks (blue bars) compared to the connection strength of observed a representative p-tau positive network from Sample 301 (red line). The structures positive for p-tau in Sample 301 had a total connection density of ∼3.5×10^−6^, which is higher than ∼98% of the possible networks that could be formed with a similar distance distribution. (E) In most brains, p-tau positive structures were more highly connected than expected by chance, as indicated by their network z scores. There was no relationship between the cluster assignments in panel A and the connectivity strength of the corresponding p-tau positve networks (compare along y-axis), or between cortical-dominant or subcortical-dominant p-tau patterns and network connection strength (compare red points and magenta points). There was no obvious similarity between the two brains that showed lower connection strength in their p-tau positive networks (308 and 371). (F) There was no relationship between the number of p-tau positive structures in a brain and network connection strength.

### Pathology co-occurs in strongly connected brain regions

Next, we tested for a relationship between regional patterns of p-tau accumulation in any given PS19 brain and underlying network connectivity, using the Allen Mouse Connectivity Atlas as the reference for normal connectivity strengths among all brain regions (*15, 33*). We compared the observed sum of connectivity strengths between all regions with detectable p-tau signal bilaterally in each individual brain to the sum of connectivity strengths derived from sets of alternate random networks of similar anatomical (Euclidean) distances (Fig. 3C, D). An individual brain’s p-tau-containing regions were more strongly connected than expected by chance (z-scores > 2; Fig. 3E, F). Notably, two brains (308 and 371) exhibited a lower connection strength among their p-tau-positive structures, which was not explained by their pathological stage or p-tau distribution pattern (Fig. 3E). We also found no correlation between the total number of p-tau positive regions with the network connectivity z-score, indicating that this relationship was not enriched in brains with higher p-tau burden (Fig. 3F). Overall, these analyses indicated that brain-wide distribution patterns of p-tau significantly correlated with their network connection strength.

### Brain networks predict patterns of pathology progression

Our previous analysis did not consider whether progression of pathology correlated with network connectivity. Thus we first identified reliable “seed regions,” defined as those with detectable p-tau in 6mo mice that were also positive for p-tau at 9-12mo. This curation produced seven consistent seed regions: anterior hypothalamic nucleus (AHN), posterior amygdalar nucleus (PA), paragigantocellular reticular nucleus, lateral part (PGRNl), Barrington’s nucleus (B), locus ceruleus (LC), entorhinal area, medial part, dorsal zone (ENTm), and medial vestibular nucleus (MV) (Fig. 4A). We used these regions as starting nodes for subsequent analyses to test the performance and validity of four models for predicting progression of tau pathology across ages: Euclidean distance between regions, (2) anterograde connectivity, (3) bi-directional connectivity (retro/anterograde), and (4) retrograde connectivity (Fig. 4B).

**Fig. 4.**
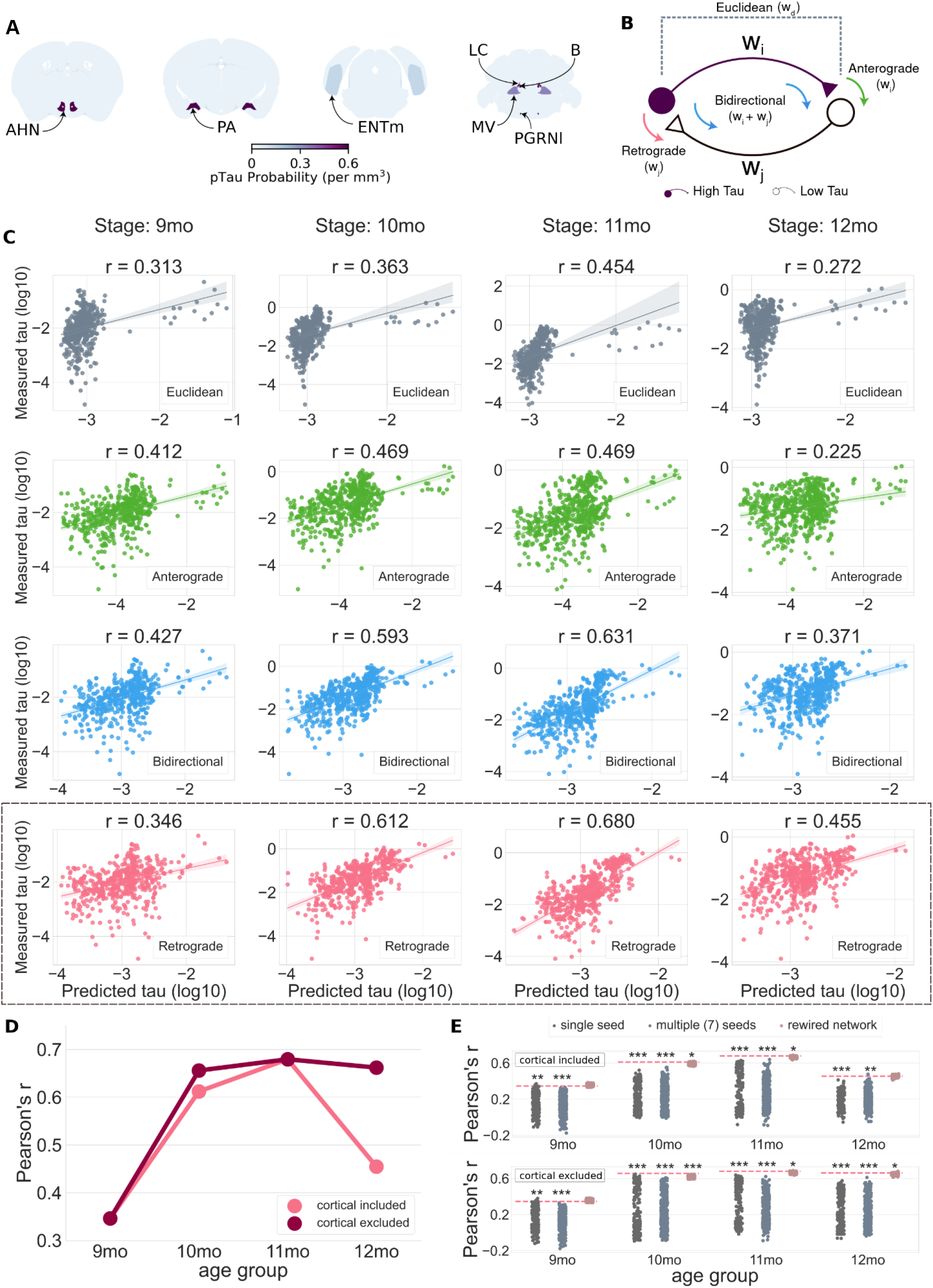
Brain networks defined with the retrograde connectome best predict spread of pathology. (A) Regions with p-tau signal in 6mo brains were identified as seed structures. Abbreviations: AHN = Anterior hypothalamic nucleus; PA = Posterior amygdala nucleus; ENTm = medial entorhinal cortex; LC = Locus ceruleus; B = Barrington’s nucleus; MV = Medial vestibular nucleus; PGRNl = Paragigantocellular reticular nucleus, lateral part. Heatmap shows the mean density of p-tau signal in each region in n=5, 6mo brains. (B) Potential mechanisms for tau spread. Neurons in structures send anterograde projections to neurons in other areas and receive projections from neurons in other areas. For simplicity, only two neurons are shown in the diagram representing cells in a region with high p-tau burden (purple) and its one neighboring structure with relatively low p-tau burden (white). If tau spreads in an anterograde direction, the amount of spread should be related to the strength of the anterograde projection (w_i) from the high p-tau region to its low p-tau neighbor(s). If tau spreads in a retrograde direction, the amount of spread should be related to the strength (w_j) of the input from the low p-tau neighbor(s) to the high p-tau region. If tau spreads in both directions, its propagation should be related to the sum of the anterograde and retrograde connection weights. If tau propagation is driven by Euclidean distance, the amount of spread should be related to the distance between regions, not the weight of their anterograde or retrograde connections. (C) Measured levels of p-tau (log scale) plotted as a function of the tau pathology predicted from propagation models based on Euclidean distance between regions (gray), anterograde connection strength (green), bidirectional connection strength (blue), and retrograde connection strength (pink). Each point represents one region from the Allen Institute CCFv3. Pearson’s r values are reported for each model-age group combination. The outliers on the right side of each scatter plot denote the seed structures. (D) Comparison of retrograde model with and without the three brains with the cortical pattern of pathology (Fig. 3A). The model predictions improved at 10 and 12mo when three brains with the cortical pathology pattern were excluded from the analysis. (E) Model performance compared to three sets of null models to evaluate specificity of seed regions and the retrograde network connections for cohorts with (top) and without (bottom) brains with the cortical pathology pattern. The null models included predictions from 315 single seeds (*i*.*e*., individual regions, dark-gray), 500 alternate combinations of 7 regions (light-gray), and 500 rewired network configurations that preserved higher-order (*i*.*e*., node degree sequence) graph statistics (pink). Each dot represents the performance of one iteration of the null model, while the horizontal dashed line specifies the performance of the proposed model that was statistically compared to the performance of the null models. Significance notation: * P<0.05, **P<0.01, ***P<0.001).

We used diffusion spread models to predict the regional distribution of p-tau pathology at later ages based on the levels of p-tau in the seed regions at 6mo. The models were applied to the brains based on age, since our previous clustering results showed that, as expected, aging was a prominent contributor to development of high levels of p-tau pathology (Fig. 3A, C4 and C5). We compared the predicted p-tau levels with the average measured p-tau per region at 9, 10, 11, and 12mo (Fig. 4C). We observed positive correlations between predicted and measured p-tau for all four models, but some outperformed others. The Euclidean distance-based model performed the worst at all ages, with the lowest Pearson’s r (Fig. 4C, gray, top row), whereas the retrograde connectivity model best predicted p-tau levels in most age groups (Fig. 4C, pink, bottom row). The anterograde connectivity model performed worse than the retrograde model in all but the 9mo age group (Fig. 4C, green). The bidirectional spread model (Fig. 4C, blue) predicted p-tau levels reasonably well and outperformed the anterograde model in every age group.

We also noted relatively lower correlations for all model predictions in the 9 and 12mo age groups. At 9mo, this may be explained by the larger time gap from the 6mo seeding epoch (3mo separation) vs. subsequent age groups (1mo separation), potentially biasing the models to better capture progression at the later ages. For 12mo, we suspected our earlier observation of two distinct deposition patterns, “Cortical” and “Subcortical”, might be a contributing factor (Fig. 3A, B). Therefore, we re-ran the same analysis without the three “Cortical” samples (2 at 12mo, 1 at 10mo). The predictive performance of the retrograde spread model indeed improved, from r=0.61 to 0.66 (10mo) and r=0.45 to 0.66 (12mo) (Fig. 4D). These results suggested the extensive cortical deposition likely reflected output from a different network pathway than that followed in most brains.

Finally, we tested whether the number and/or the specific set of seed regions impacted performance of the spread models. We compared the Pearson’s r values derived from the retrograde spread model against the performance of three different “null” model predictions based on: (1) a single seed region randomly selected from the entire set of summary structures, randomized sets of n=7 seed regions from the entire set of summary structures, and (3) “rewired,” or plausible alternative sets of connectivity strengths (*34*) between the n=7 p-tau seed regions (Fig. 4E). Using non-parametric permutation testing, we found the retrograde spread model with the curated set of seven seed regions significantly (p<0.01) outperformed both single (dark-gray) and multiple seed (light gray) null models in all age groups regardless of cortical mice inclusion. The retrograde model also significantly (p<0.05) outperformed the “rewired” network null models at 10 and 11mo with inclusion of cortical brains, and in 10, 11, and 12mo age groups with the three cortical brains excluded. These results further indicated that p-tau spread in PS19 mice was driven specifically from these seven seed regions and was strongly related to the underlying connectome architecture mapped in the Allen Mouse Brain Connectivity Atlas. Overall, these data indicated that neuronal p-tau accumulation patterns across age were primarily propagated via the anatomical connectome as opposed to absolute Euclidean proximity between the regions, and that spread between regions was biased towards the retrograde direction.

## Discussion

It has remained unknown, but of critical importance, whether the axonal projections that comprise brain networks mediate propagation of spontaneous tau pathology, both for an understanding of disease mechanisms and for preclinical translational studies in mouse models.

### Mapping whole brain tau pathology in 3D

We used a novel staining protocol with a fluorescently conjugated camelid anti-p-tau nanobody and STPT to generate high resolution whole brain images of p-tau accumulation in PS19 mice aged 6-12mos. The use of nanobodies, which can in theory be engineered to bind any substrate, to map proteins at subcellular resolution is a powerful approach with potentially broad applications. Because VHH-A2 was directly conjugated to AlexaFluor 488, whole-brain staining of fixed, permeabilized mouse brains was performed in a single, highly efficient step. This method, combined with STPT, reduces some barriers to whole brain mapping using alternative methods such as traditional histological sectioning or lightsheet microscopy. It was also enabled by our development of an informatics pipeline incorporating supervised machine learning to detect and quantify cellular p-tau pathology, with registration of the acquired image stacks to a standard 3D anatomical reference space—the Allen CCFv3 atlas. We anticipate that this approach could be used to study many aspects of brain biology and neuropathology.

### Progressive tau pathology follows brain networks

Propagation of tau pathology based on prion mechanisms requires that tau protein be expressed in a native, “seedable” state in vulnerable cells, with the progressive aggregation mediated by trans-cellular movement of seed-competent tau. Although proposed by many, it has remained unclear whether spontaneously occurring tau pathology in a human, or in a mouse model, spreads via brain networks. In humans, studies of the role of network involvement have combined functional connectivity or tractography, with imaging of tau pathology based on positron emission tomography, or, as a surrogate, brain atrophy (*7*). Each of these methods, while state-of-the-art in humans, has poor cellular resolution, orders of magnitude below what was achieved here via STPT. In mouse models, network studies have been based on inoculation of seeds (*9, 12*), which does not reflect a normal disease process, and is prone to experimental artifact.

The Allen Mouse Connectivity Atlas and CCFv3 resources facilitated our analyses of brain-wide tau propagation patterns across discrete anatomical regions (*15*). By registering whole brain images of p-tau deposition in PS19 mice into the annotated Allen CCFv3, similar to recent studies (*13, 22*), we identified seven regions which consistently exhibited pathology at 6mo and all later ages. We determined that the emergence of additional sites of pathology was related to age, but not exclusively. The areas of the brain with tau pathology in any given animal exhibited stronger network inter-connectivity than would be expected from a random set of regions. We also found that the connectivity strengths between regions in the retrograde direction best predicted progression of p-tau levels with increasing age. In contrast, anatomic proximity, with a few exceptions, had relatively weak predictive power. Thus, the simplest interpretation of our data is that pathology begins in discrete areas in PS19 mice, before propagating via network connections with a retrograde bias.

We made a surprising observation of two relatively discrete patterns of late stage pathology: cortical vs. subcortical. In the three cases with dominant cortical pathology, selected seed regions did not effectively predict their involvement in later ages, reducing the apparent power of the network model. However, when the three mice with cortical involvement were not included in the analysis, the predictive power improved. The origins of these two distinct late stage pathological patterns in isogenic PS19 mice are not clear. They could arise from variation in earliest regions of initiation, differences in neural development, strain composition, or other unknown factors that impact the emergence of tau pathology.

It is not yet technically possible to track in real time the physical propagation of protein pathology across distributed brain networks *in vivo, i*.*e*., transcellular movement of tau seeds. However, our observation that emergent cellular pathology in a mouse model with pan-neuronal mutant tau expression is accurately predicted by a retrograde propagation model suggests that in mice, as has been long hypothesized in humans (*35*), spontaneous tau prion phenomena underlie neuropathological progression. It also indicates that the PS19, and perhaps other endogenous tauopathy models, remain useful for analysis of mechanisms of propagation, and development of diagnostic and therapeutic strategies that can be translated to humans.

## Supporting information

All supplementary materials

Visualization of phospho-tau accumulation patterns in three major clusters of brains

## Acknowledgments

All TissueCyte imaging and quantification was performed in the UT Southwestern Whole Brain Microscopy Facility (RRID:SCR_017949). We thank the Allen Institute founder, Paul G. Allen, for his vision, encouragement, and support. The project described was supported in part by awards from the National Institutes of Health to MID, JPM and JAH. Its contents are solely the responsibility of the authors and does not necessarily represent the official views of the National Institutes of Health.

## Author contributions

Conceptualization: MID, JPM, JAH, NB, JDW, DMOR

Methodology: MID, JPM, JAH, NB, JDW, DMOR, TLT, ADA, AN, BD, SB, JEK, PL

Resources: BD, SB, PL, JEK, MID

Investigation: DMOR, JDW, NB, TLT, JPM, JAH

Visualization: DMOR, JDW, NB, ADA, AN, JPM, JAH

Funding acquisition: MID, JPM, JAH

Project administration: MID, JPM, JAH, DMOR

Supervision: MID, JPM, JAH

Writing – original draft: DMOR, JDW, NB, JAH, JPM, MID

Writing – review & editing: DMOR, JDW, NB, JAH, JPM, MID, ADA, AN

### Data and materials availability

Registered 3D image stacks of p-tau probability maps, the matrix of p-tau intensity per CCFv3 annotated brain region for each brain, and associated custom code used in visualization and analyses are available at https://doi.org/10.5281/zenodo.7942188.

## Supplementary Materials

Materials and Methods

Figs. S1 to S5

Table S1

References (*36–40*)

Movie S1

