## Supplementary material for "Endogenous pathology in tauopathy mice progresses via brain networks": All supplementary materials

### Materials and Methods

Animals

PS19 mice expressing 4R1N P301S human tau under the murine prion promoter (*23*) and mice lacking endogenous tau (TauKO) (*31*) at ages from 3-12 months were used. The mice used in this study included both males and females, with the exception of the 6mo group of PS19 mice which were all female (Table S1). All mice involved in this study were housed under a 12-h light/dark cycle and were provided food and water ad libitum. All experiments involving animals were approved by the University of Texas Southwestern Medical Center Institutional Animal Care and Use Committee (IACUC).

Tissue harvest and immunostaining

PS19 animals were sacrificed at 3 (n=5), 6 (n=5), 9 (n=3), 10 (n=6), 11 (n=5), or 12 (n=4) months of age via transcardial perfusion with PBS followed by 4% paraformaldehyde (PFA; w/v) in PBS. Three TauKO mice at 12 or 14 months of age were additionally sacrificed and perfused in the same manner. Brains were extracted and post-fixed overnight in 4% PFA at $4^{o}$C. Brains were rinsed in PBS before staining and permeabilized in 10ml of 0.2% Triton-X-100 (Surfact-Amps X-100; ThermoFisher #28314) + PBS for 48hrs at room temperature (RT) while gently shaking, with three total washes. Brains were incubated in 162.5μg/ml VHH-A2 previously conjugated to AlexaFluor 488 (VHH-A2-488; a nanobody which specifically recognizes pathological tau phosphorylated at S422) (*16*), diluted in PBS+0.2% (v/v) Triton X-100 to a total volume of 1ml for 6 days at $4^{o}$C with constant rocking. Nanobody solution was removed and brains were washed in 0.2% Triton-X-100 + PBS for 2 days at $4^{o}$C with gentle shaking and solution was changed twice a day. Brains were then rinsed well in PBS in preparation for agarose embedding as described below.

Agarose embedding

A modified agarose embedding method was employed for the majority of the brains used in this study, which incorporated infiltration of the immunostained brains with acrylamide prior to agarose embedding. The brains were soaked overnight at 4°C in a solution of Surecast (ThermoFisher #HC2040), with a total concentration of 4% Surecast and 0.5% VA-044 activator, in excess volume (5mL per brain), diluted in Phosphate Buffer (PB; 0.42g/L Monobasic sodium phosphate, 0.92g/L Dibasic sodium phosphate). The brains were then allowed to equilibrate to RT for 1 hour while gently shaking. Each brain was transferred to a cryomold (VWR, #15160-215), which was filled with the prepared acrylamide solution. The mold was covered securely with foil and incubated in an oven at 40°C for two hours. After polymerization was complete, the excess acrylamide was carefully removed from around the brain. The acrylamide-infused tissue was then embedded in oxidized agarose for vibratome sectioning and TissueCyte imaging as previously described (*17-18*,*20*). A subset of brains was embedded in agarose but did not receive the pre-treatment with acrylamide using an earlier version of the embedding protocol.

Serial Two-Photon Tomography (STPT)

STPT was performed on the TissueCyte 1000 (TissueVision, Inc., Newton, MA) instrument essentially as described (*17-18, 20*). The agarose blocks containing the brain samples were attached to a custom magnetic slide with superglue and placed on a magnetized stage within an imaging chamber filled with PB. STPT imaging, a block-face technique, was used to acquire a series of 2-dimensional (2-D) mosaic images in the coronal plane (*19*). For this study, three optical planes were imaged at 25, 50, and 75 µm below the cut surface, followed by a vibrating microtome cut at 75 µm (blade vibration frequency of 70 Hz, advancement velocity 0.5 mm/s). The excitation laser (MaiTai DeepSee, SpectraPhysics/Newport, Santa Clara, CA) wavelength was tuned to 800 nm to efficiently excite AlexaFluor 488. Fluorescent signals from three emission channels were collected with a predetermined photomultiplier tube voltage of 720 V using appropriate emission filters encompassing red (>560nm), green (500-560nm), and blue (<500nm) wavelengths. This process produced 190 physical sections and 570 2-D stitched coronal section images (9 by 13 mosaic collected for each coronal section) with a lateral resolution of 0.875 µm/pixel and axial resolution of 25 µm, generating ~280 gigabytes of raw data per brain.

Image processing

Raw image tiles were trimmed and subjected to flat field correction, then stitched into 2-D mosaic coronal section images using Autostitcher software (TissueVision, Inc.). Using custom written macros (MATLAB), 3-channel merged, contrast adjusted, maximum intensity projection (MIP) images were generated for each physical section (using the 3 optical planes collected for each physical section). The merged MIP images were downsampled to 1.5µm/px, and contrast was linearly adjusted to match intensity histograms as much as possible to support the subsequent analysis pipeline. A manual quality control process was employed for each immunostained brain to eliminate samples with obvious structural flaws or imaging artifacts. Brains showing excessive vascular autofluorescence signal, no identifiable neuronal phospho-tau staining, or missing portions of the images due to sectioning or dissection artifacts were excluded from analysis.

Automated image quantification

Our pipeline for quantification of STPT whole brain image datasets was previously described in detail elsewhere (*17-18, 20*). Briefly, the pipeline consists of supervised machine learning based pixel classification, registration of the brain volumes into the reference atlas (Allen Institute CCFv3.0; (*21*)) and automated quantification of fluorescent signals in all annotated brain regions. Supervised machine learning-based pixel classification was performed using a pixel-wise random forest (RF) classifier in ilastik ("Pixel Classification" applet; (*36*)), which allowed isolation of fluorescent signals of interest (e.g., VHH-stained tau deposits) across brain regions that had variable fluorescent backgrounds. A subset of images from each brain was included in the RF model training set (2-3 images per brain). A single RF classifier was trained and used for all brains in the dataset. Probability maps for were exported as scaled 8-bit images (a pixel intensity value of 255 equated to a 100% probability that a pixel belonged to a signal label). Probability maps were subjected to median filtering prior to analysis to exclude high contrast noise artifacts. Registration to the CCFv3.0 was performed using the raw red (>560 nm) channel autofluorescence signal using a combination of translation, affine, and bspline transformations with SimpleElastix (*37*). The registration transformation parameters for each brain were applied to the probability map datasets, resulting in registered probability maps of phospho-tau deposition. Finally, the normalized intensity of all voxels lying within CCFv3.0 annotated regions was quantified using custom MATLAB scripts, producing a matrix of phospho-tau signal intensity for each sample, brain region, and hemisphere.

Background subtraction

Our automated classifier was not always able to differentiate between true positive signal (Fig. S5A) and autofluorescence from lipofuscin or other background fluorescence in the tissue (Fig. S5B), but true positives and false positives could be differentiated by eye. To remove the automatically detected false positive values in our per-structure quantification, we took advantage of two tau KO brains that were processed alongside our experimental samples as negative controls. Brains from five additional 3-month old PS19 mice – an age with minimal tau pathology (*23*) – were immunostained and imaged using the same methods to assist in the determination of background staining in PS19 brains (data not shown). We also manually scored a subset of regions in twelve brains. We visually checked 17-112 structures per brain in all 3- and 6-month-old brains plus three 12-month-old brains (1019 structures total). Since fewer than 10% of the structures in the 3-month-old brains were true positives (36/336) and most of this signal was extremely sparse, we included the 3-month-old brains with the tau KO controls and defined a threshold using the maximum value per structure in the 3-month-old and tau KO brains, excluding the values for the 36 true positive structures. We then subtracted these threshold values from the quantified density per structure in all brains. The distribution of density values for all true-positive and true-negative structures is plotted in Fig. S5C. This per-structure background subtraction eliminated 488 false positives and resulted in 70 true positive structures falling below the threshold and being assigned a zero value (Fig. S5D). The whole brain p-tau density for each brain is shown in Fig. S5E before and after baseline subtraction.

Regional distribution analysis

To test whether the distribution of tau pathology in each brain was related to structural connectivity, we took advantage of a brain-wide model that converts the density of axonal projections measured by stereotaxic injections in the Allen Mouse Brain Connectivity atlas (*15*) into a structure-wise connectivity weight for all annotated structures in the common coordinate framework ((*33*) regionalized voxel model, normalized connection density). For each brain, we first calculated the sum of the connectivity weights for all structures containing tau pathology, then compared it to a distribution of randomly sampled networks of the same size with similar inter-regional distances. The procedure for this selection is as follows:

Given a set of N structures containing tau pathology:

1) Randomly draw a set of N structures

2) Compute the pairwise inter-regional distances for the set of sampled regions.

3) Compute the Kolmogorov-Smirnov statistic to measure the difference in distributions of distances for the sampled and tau correlated networks

a) If the KS statistic shows a significant difference in distributions (having a p-value < 0.01), reject the sample and return to (1)

b) Else, return the sample.

The above procedure was repeated 1000 times, after which the normalized connection density of the tau network was compared to the normalized connection density of the random sample networks. Since these connectivity measures are log-normally distributed (*15*, *33*), the t statistic is computed in log-transformed space to test the significance of the difference.

Computational models of pathology spread

Network diffusion models have been used to characterize and predict pathological protein spread patterns implicated in Alzheimer's and Parkinson's disease in mice and humans (*13, 38-40*). These models hypothesize linear diffusion of pathology over the brain connectome composed of the white-matter tracts. Thus, the pathology spread from a given brain region to its neighbors is a function of their 1) strength of connections and 2) pathology differentials. In this work, we applied the network diffusion model initialized with multiple seed structures on a directed brain graph to predict regional spread patterns of p-tau with respect to age.

To identify seed structures for the network diffusion model, we included regions that were manually validated as true positives in all the 6-month-old brains and were informatically identified as true positives at all subsequent ages. We then included only structures in the top 5% of all quantified values at 6 months for either total tau or tau density per structure. Only seven structures met all of these criteria: AHN, PA, PGRNl, B, LC, ENTm, and MV' (see Fig. 4A).

We constructed the directed brain graph $G = \{V, E\}$ with $V$ nodes (gray matter regions) and $E$ edges (white matter projections). The nodes ($\mathrm{N_v} = 625$) comprise 310 bilateral and 5 medial cortical and subcortical gray matter regions defined in the Allen Reference Atlas ontology (*i.e.* CCFv3 "summary structures" - see Fig. 3). The edge strengths were quantified using the green fluorescent protein measures from the Allen Mouse Brain Connectivity Atlas (*15*).

We represent the adjacency matrix of $G$ as $\mathrm{A} = [\mathrm{A_{ij}}]$, where $\mathrm{i}'$th row contains connection strengths from the region $i$ to all other ipsi-, medial, and contra-lateral brain regions.

We denote the measured pathology burden of all nodes at a given time epoch (i.e. mouse age relative to the seed time point) as a vector $\mathrm{x(t)}$. We initialize the model with seed structures at the 6 month time point and then compute the predicted tau burdens, $\hat{x}(t)$ at subsequent $t = 3,4,5,6$ month-intervals with the following equation:

$$\hat{x}(t) = e^{-\beta L t} x(0)$$

and,

$$L = D - A$$

where, $L$ is a graph Laplacian matrix calculated from the graph out-degree matrix $D$ (i.e. sum of outgoing connection strengths for each node) and the adjacency matrix $A$. Hyperparameter $\beta$ is a diffusion rate constant which scales the global speed of spread for a given network comprising nodes and edges quantified in arbitrary and relative units of pathology and projection strengths, respectively.

We evaluated the model performance with Pearson correlation coefficient between $\log\left(\hat{x}(t)\right)$ (predicted) and $\log\left(x(\mathrm{t})\right)$ (measured) pTau levels at each age. The optimal model was identified by fitting model instances for a range of $\beta$ values ($\beta \in [1\mathrm{e}^{-5}, 100]$) and selecting the one that maximized the average Pearson correlation coefficient over all time points. We checked that the selected $\beta$ remained well within the predefined bounds for each fitted model, and the model performance remained stable over small perturbations of $\beta$. With this setup, we carried out two experiments with the seven identified seed structures.

In the first experiment, we compared the model performance across three anatomical adjacency matrices and a single Euclidean adjacency matrix (see Fig. 4B,C). The anatomical adjacency was derived using 1) anterograde 2) retrograde, and 3) bidirectional projection strengths. The retrograde adjacency matrix was defined as the transpose of the anterograde connectivity matrix. The bidirectional adjacency matrix was calculated as the sum of anterograde and retrograde connectivity matrices. Euclidean adjacency was calculated as the inverse Euclidean distance between the center-of-mass vertices of anatomical regions. Four separate models were fitted for each of these adjacency measures.

In the second experiment, we compared the retrograde model performance with and without three mice with substantial levels of cortical p-tau (see Fig. 4D). This involved refitting the model with exclusion of one brain at 10mo and two brains at 12mo.

Finally, we fitted three sets of null models to evaluate retrograde model performance against the specificity of seed structures as well as the network configuration using non-parametric permutation tests (see Fig. 4E). In the first set, we fitted separate models with each region from the entire set of bilateral structures set as a single seed. In the second set, we evaluated models with 500 alternate combinations of seven seed regions. Both of these sets of models estimate the null performance distributions with single and multiple random seed initializations. Lastly, in the third set, we evaluated performance of 500 rewired network configurations that preserve higher-order (*i.e.* node degree sequence) graph properties (*34*). The network null models test whether the performance is driven by the topology of the anatomical connectome represented by A or if it could be obtained with alternative graphs with similar higher-order graph statistics. The p-values were calculated as the percentage of times null models yielded better performance than the retrograde model at each age.

**HJ8.5 Immunostaining and quantification of total tau levels**

PS19 animals at 3, 6, 9 and 12 months of age were subjected to transcardial perfusion with PBS and 4% PFA. Brains were extracted and postfixed in 4% PFA overnight at 4$^{o}$C and transferred to 30% sucrose (w/v) in PBS until sunk. Coronal sections were prepared on a freezing microtome at 50 μm thickness. Selected sections spanning the rostrocaudal extent of the hippocampus were immunostained with HJ8.5 monoclonal antibodies (*10*) to detect expression of total human tau. Briefly, free-floating sections were permeabilized for 15 min at RT in PBS+0.2% (v/v) Triton-X100 and blocked in NGS blocking solution (10% normal goat serum, 5% (w/v) BSA, 0.2% Triton X-100 in PBS) for at least 1 hour at RT. The sections were incubated with HJ8.5 antibodies sourced from the Diamond laboratory overnight in well plates at 4$^{o}$C diluted in NGS blocking solution. Sections were washed 3 times 10 min each in 0.025% Triton X-100 in PBS at RT, followed by incubation with goat-anti-mouse AlexaFluor 594 secondary antibodies (1:500; ThermoFisher #A-11032) in NGS blocking solution for 1 hour at RT in the dark. Sections were washed 3 times 10 min each in 0.025% Triton X-100 in PBS at RT, stained with DAPI and coverslipped. Whole slide images of the red (HJ8.5) and blue (DAPI) channels were acquired at 20X magnification on the Zeiss Axioscan.Z1. The measurement tool in Zeiss Zen Lite Blue was used to obtain intensity measurements for HJ8.5 staining in the red channel for each section from each animal. Intensity results from sections stained with primary HJ8.5 antibodies were normalized to section area and the corresponding no primary control section from each animal was used for background subtraction.

**AT8 Immunostaining**

Selected 75μm sections from brains that had been subjected to whole brain VHH-A2-488 staining and STPT were recovered after imaging and stained with AT8 primary antibodies to assess the overlap of p-tau staining between the two antibodies. Briefly, free-floating sections were permeabilized for 15 min at RT in PBS+0.2% (v/v) Triton-X100 and blocked in NGS blocking solution (10% normal goat serum, 5% (w/v) BSA, 0.2% Triton X-100 in PBS) for at least 1 hour at RT. The sections were incubated with biotinylated AT8 antibodies (AT8b, Fisher Scientific #ENMN1020B) overnight in a well plate at 4$^{o}$C diluted 1:500 in NGS blocking solution. Sections were washed 3 times 10 min each in 0.025% Triton X-100 in PBS at RT, followed by incubation with AlexaFluor 594-streptavidin (1:500; ThermoFisher #S11227) in NGS blocking solution for 1 hour at RT in the dark. Sections were washed 3 times 10 min each in 0.025% Triton X-100 in PBS at RT, stained with DAPI and coverslipped. Whole slide images of the blue (DAPI), green (VHH-A2-488) and red (AT8) were acquired at 20X magnification on the Zeiss Axioscan.Z1.

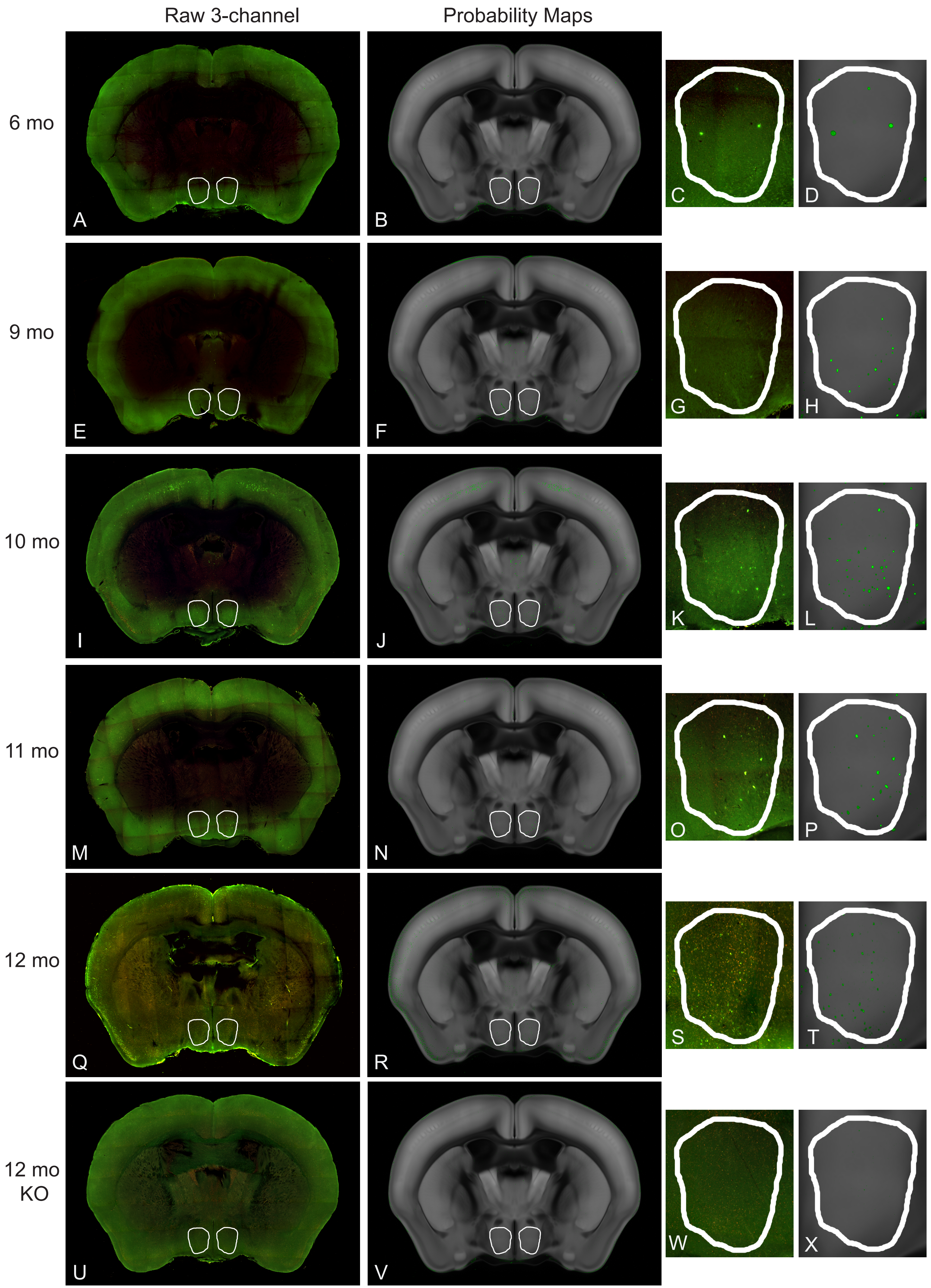

Raw 3-channel
Probability Maps
6 mo
9 mo
10 mo
11 mo
12 mo
12 mo KO
A
B
C
D
E
F
G
H
I
J
K
L
M
N
O
P
Q
R
S
T
U
V
W
X

**Fig. S1. Fluorescent immunostaining of p-tau using VHH-A2-488 in PS19 animals at 6, 9, 10, 11, and 12 months of age and a TauKO mouse at 12 months of age.** Intact mouse brains were subjected to immunostaining with VHH-A2-488 and imaged using serial two-photon tomography. Panels A, E, I, M, Q, and U present representative raw fluorescence images from a single brain for each time point at the level of the anterior hypothalamic nucleus (AHN), a seed region showing consistent pathology at all mouse ages used in this study. AHN is annotated in white in all images. Fluorescent p-tau staining is shown in green. The red channel represents tissue autofluorescence. Panels B, F, J, N, R, and V show the corresponding probability maps of segmented p-tau signal (green) overlaid onto the CCFv3 average template (grey). Panels C, G, K, O, S, and W show enlargements of the raw fluorescence from right AHN from each brain, and panels D, H, L, P, T, and X show the corresponding probability maps. Positive p-tau staining in the AHN is apparent in each of the PS19 brains at different ages and was accurately segmented using our classifier, but no specific staining was observed in 12 month old TauKO brains.

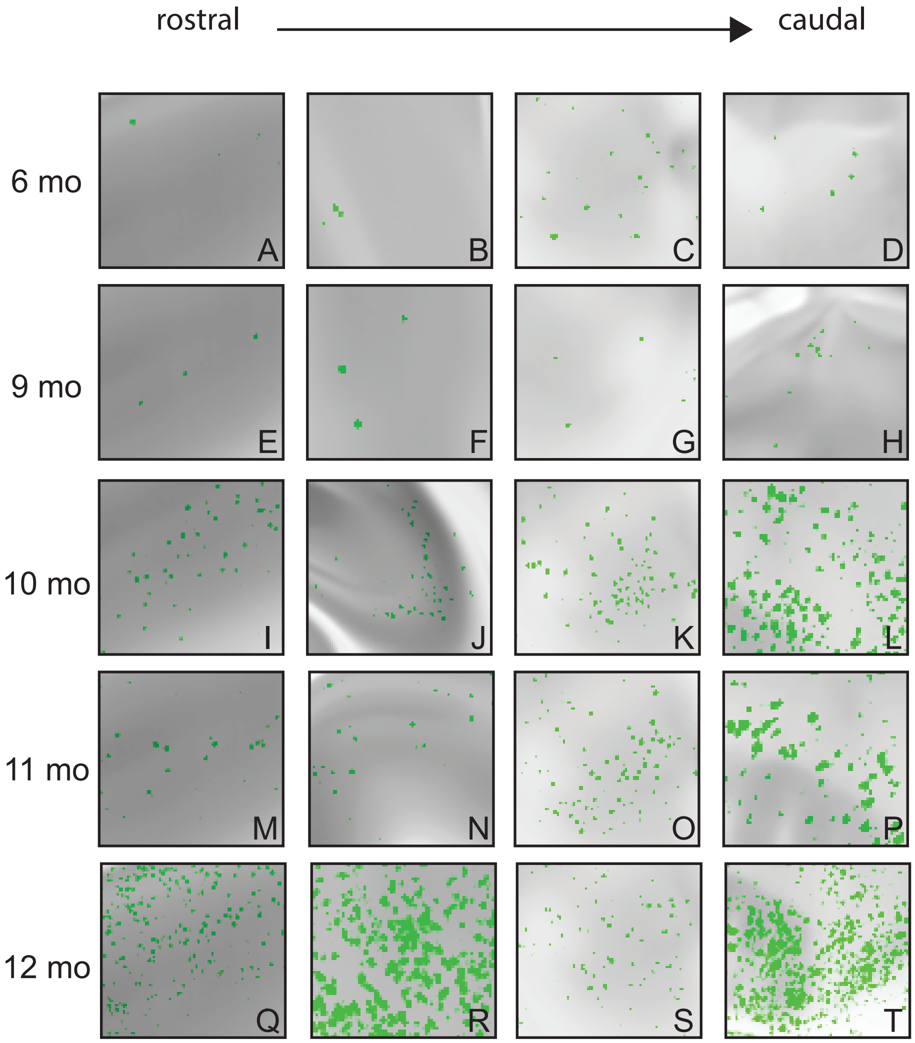
rostral
caudal
6 mo
9 mo
10 mo
11 mo
12 mo
A
B
C
D
E
F
G
H
I
J
K
L
M
N
O
P
Q
R
S
T

**Fig. S2. Phospho-tau pathology initiates in brainstem and other caudal brain regions and progressively accumulates through the brain towards rostral areas.** All panels show high resolution views of areas indicated by asterisks (*) in Fig. 2. Maximum intensity projections (MIP) of p-tau probability maps for each timepoint at selected anatomical levels are shown. Images are extracted from the four coronal sections spanning the rostro-caudal extent of the brain at the levels of the anterior cortex, dorsal hippocampus, midbrain and cerebellum as shown in Fig. 2 for brains of the indicated ages (6mo = Panels A-D, 9mo = Panels E-H, 10mo = Panels I-L, 11mo = Panels M-P, 12mo = Panels Q-T)**.** P-tau probabilities are shown in green and the CCFv3 average template in grey. The total numbers of brains in the dataset at each timepoint (6 mo = 5, 9 mo = 3, 10 mo = 6, 11 mo = 5, 12 mo = 4) were used to generate MIP images of each physical section. Progressive accumulation of p-tau is visible across many brain regions, notably brainstem, hippocampus and ventral cortical areas including entorhinal (ENT), cortical amygdalar (COA) and piriform (PIR).

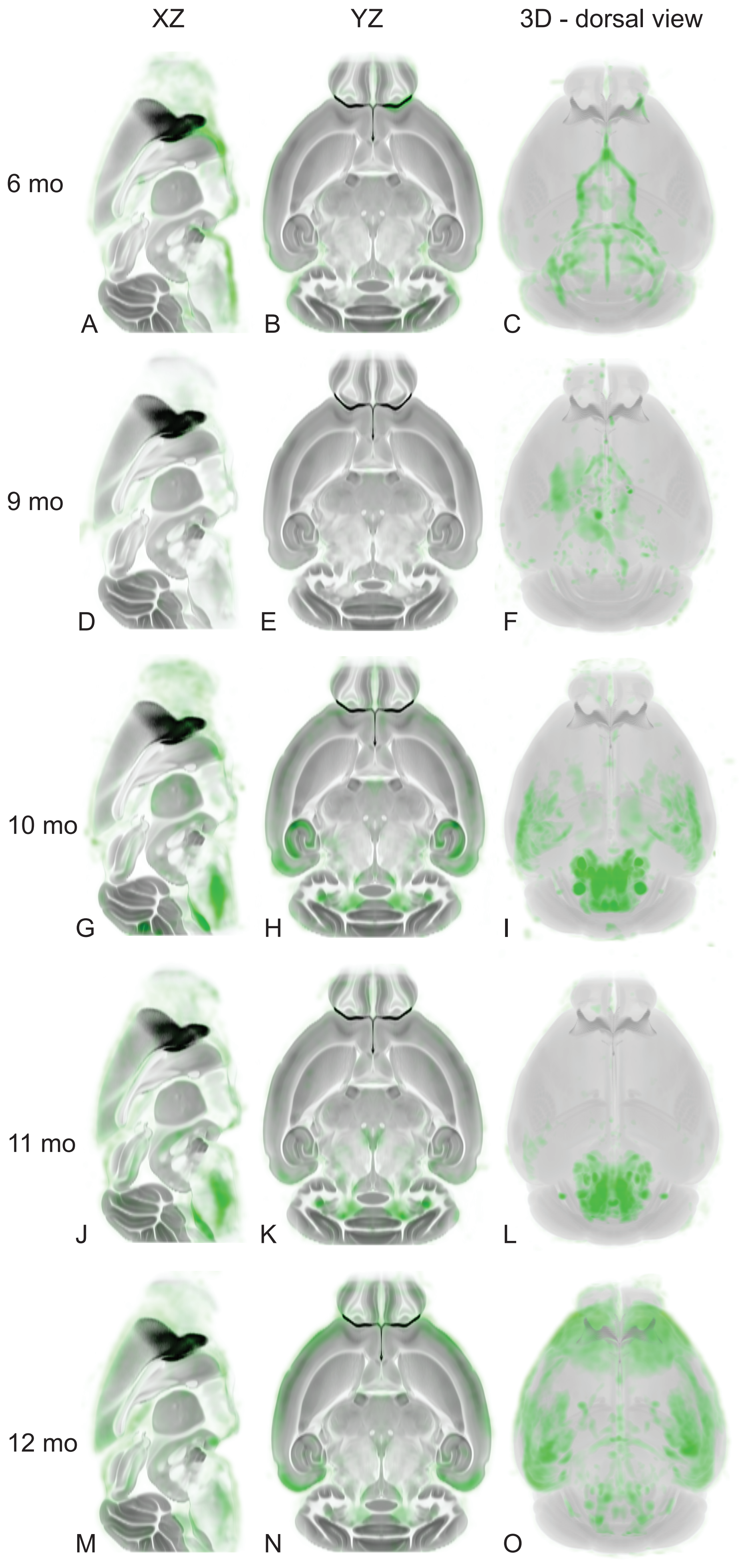

XZ
YZ
3D - dorsal view
6 mo
9 mo
10 mo
11 mo
12 mo
A
B
C
D
E
F
G
H
I
J
K
L
M
N
O

**Fig. S3. Average brainwide p-tau distribution in PS19 mice at each timepoint.** Registered p-tau probabilities from multiple brains at each timepoint (6, 9, 10, 11, and 12 months) were averaged. P-tau probabilities are shown in green and the CCF average template in grey. Orthogonal XZ (Panels A, D, G, J, M) and YZ (Panels B, E, H, K, N) planes are shown, as well as 3D renderings of the whole average brain from the dorsal view (Panels C, F, I, L and O). Number of brains averaged for each timepoint: 6mo = 5, 9mo = 3, 10mo = 6, 11mo = 5, 12mo =4. Progressive accumulation of p-tau is visible across many brain regions, notably brainstem, hippocampus and cortex.

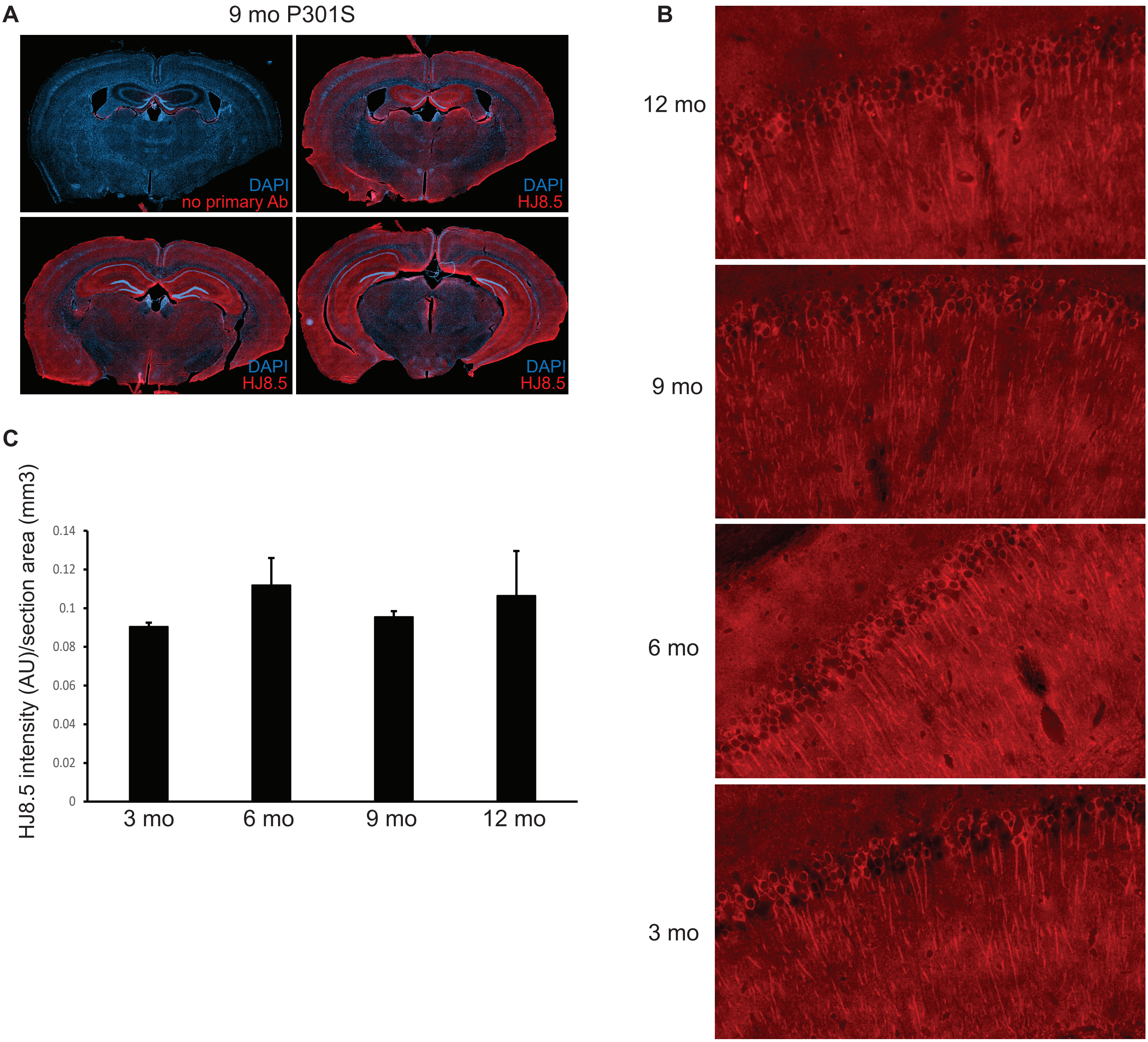
A
9 mo P301S
DAPI
no primary Ab
DAPI
HJ8.5
DAPI
HJ8.5
DAPI
HJ8.5
B
12 mo
9 mo
6 mo
3 mo
C
HJ8.5 intensity (AU)/section area (mm3)
0.14
0.12
0.1
0.08
0.06
0.04
0.02
0
3 mo
6 mo
9 mo
12 mo

**Fig. S4. Total levels of human tau do not vary with age in PS19 mice.** Brains were harvested at 3, 6, 9, and 12 months and serial frozen sections were subjected to immunostaining with HJ8.5 antibodies to label total human tau. A) Representative whole section images of the series of three sections at different anatomical levels from a 9 month old brain stained with HJ8.5 (Top right and bottom 2 panels) or a no primary control section from the same brain (top left panel). HJ8.5 staining is shown in red and DAPI nuclear staining is shown in blue. B) Full resolution (20x) images of hippocampal area CA1 from brains at each of the indicated ages showing robust HJ8.5 staining in red. C) Quantification of HJ8.5 staining intensity across six serial sections for each age group (one series of three sections at anatomical levels as shown in Fig. S4A from two brains per age group). Error bars = SEM. No significant differences were observed in total tau levels between ages.

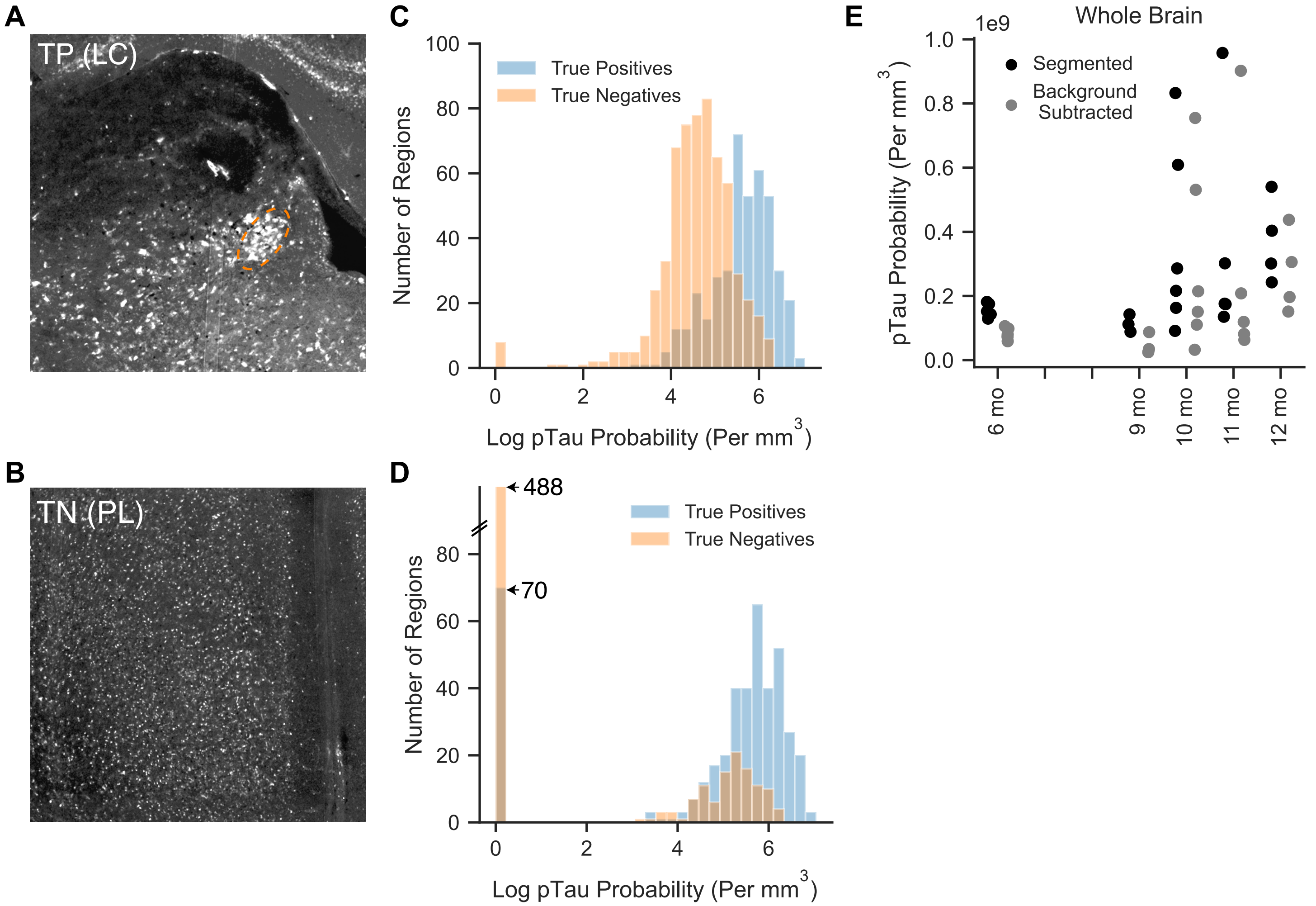

A
TP (LC)
B
TN (PL)
C
True Positives
True Negatives
Number of Regions
Log pTau Probability (Per mm^3)
D
488
70
True Positives
True Negatives
Number of Regions
Log pTau Probability (Per mm^3)
E
Whole Brain
1e9
Segmented
Background Subtracted
pTau Probability (Per mm^3)
6 mo
9 mo
10 mo
11 mo
12 mo

**Fig. S5. Development of background subtraction method to reduce contributions of automatically detected false positive phospho-tau staining to analysis.** Example images showing true positive p-tau immunostaining in locus ceruleus (LC, A) and true negative background immunostaining in the prelimbic area (PL, B). Dashed oval in panel A shows approximate LC boundaries. C) Histogram showing the probability values for regions scored as True Positives (blue) and True Negatives (Orange). D) Histogram showing probability values for the same regions after background subtraction. E) Whole brain p-tau probabilities before (black) and after (gray) background subtraction.

| Sample ID | Age (mo) | Sex | Cluster |
|---|---|---|---|
| 297 | 12 | F | 5 |
| 298 | 12 | F | 4 |
| 296 | 12 | M | 4 |
| 371 | 12 | M | 5 |
| 299 | 11 | F | 2 |
| 300 | 11 | F | 2 |
| 301 | 11 | M | 4 |
| 620 | 11 | M | 4 |
| 625 | 11 | M | 2 |
| 711 | 10 | F | 5 |
| 716 | 10 | F | 3 |
| 717 | 10 | F | 2 |
| 715 | 10 | M | 3 |
| 719 | 10 | M | 4 |
| 721 | 10 | M | 3 |
| 308 | 9 | F | 1 |
| 309 | 9 | F | 3 |
| 710 | 9 | M | 2 |
| 748 | 6 | F | 1 |
| 749 | 6 | F | 1 |
| 750 | 6 | F | 1 |
| 751 | 6 | F | 1 |
| 752 | 6 | F | 1 |

**Table S1. Sex and phospho-tau cluster assignment information for PS19 mice analyzed in this study.** Within the limitations of our study that was not powered to detect sex differences as a primary outcome, sex did not appear to affect either the pathological tau burden or its accumulation pattern at any age.

**Movie S1. Visualization of phospho-tau accumulation patterns in three major clusters of brains.**

Phospho-tau probabilities were averaged for 9 brains comprising clusters 3, 4, and 5 as shown in Fig. 3 which displayed either subcortical or cortical patterns of pathological tau accumulation in our network connectivity analysis. Images are presented as an overlay with each of 190 coronal sections displayed in sequence from caudal to rostral. Red = subcortical pattern with heaviest tau burden (cluster 4 brains 301, 620 and 719), green = cortical pattern (cluster 5 brains 297, 371 and 711), blue = subcortical with moderate tau burden (cluster 3 brains 715 and 721 and cluster 4 brain 298). The overlapping nature of the less affected (blue) and more highly affected (red) subcortical pattern brains is apparent, with additional regions showing tau accumulation in the brains with overall heavier pathological tau burden. The divergent pattern for the subset of brains with cortical accumulation compared to subcortical is also evident.
